# Outdoor Atmospheric Microbial Diversity is Associated with Urban Landscape Structure and Differs from Indoor-Transit Systems as Revealed by Mobile Monitoring and Three-Dimensional Spatial Analysis

**DOI:** 10.1101/2020.06.17.157651

**Authors:** JD Stewart, P. Kremer, K.M. Shakya, M. Conway, A. Saad

## Abstract

Microbes are abundant inhabitants of the near-surface atmosphere in urban areas. The distribution of microbial communities may benefit or hinder human wellbeing and ecosystem function. Surveys of airborne microbial diversity are uncommon in both natural and built environments and those that investigate diversity are stationary in the city, thus missing continuous exposure to microbes that covary with three-dimensional urban structure. Individuals in cities are generally mobile and would be exposed to diverse urban structures outdoors and within indoor-transit systems in a day. We used mobile monitoring of microbial diversity and geographic information system spatial analysis, across Philadelphia, Pennsylvania, USA in outdoor and indoor-transit (subways and train cars) environments. This study identifies to the role of the three-dimensional urban landscape in structuring atmospheric microbiomes and employs mobile monitoring over ~1920 kilometers to measure continuous biodiversity. We found more diverse communities outdoors that significantly differ from indoor-transit air in microbial community structure, function, likely source environment, and potentially pathogenic fraction of the community. Variation in the structure of the urban landscape was associated with diversity and function of the near-surface atmospheric microbiome in outdoor samples.

**Importance:** Global nutrient cycling and human health depend on the rich biodiversity of microorganisms. The influence of the urban environment on microbiomes remains poorly described despite cities being the fastest growing ecosystems. All life is exposed to the atmosphere and thus discerning what microbes are present and what their functions may be is critical to create resistant and resilient cities under climate change. This study combines a spatially explicit analysis of urban structure with mobile monitoring of the atmospheric microbiome.

## Introduction

Microbes are abundant inhabitants of the near-surface atmosphere (Burrows et al., 2009) whose presence and distributions in urban areas have implications on public health and ecosystem function. The atmosphere serves as both a source and a sink for microorganisms that may return to the surface through dry or wet deposition (Chate et al., 2003; Wu et al., 2018). Furthermore, urban areas are the fastest growing ecosystems (Grimm et al., 2008) and are composed of heterogeneous landscapes (Cadenasso et al., 2007) with diverse land covers combined in various configurations and proportions. Many studies of urban structure rely on remotely sensed landcover datasets that distinguish between the built and natural elements of the landscape; however, they ignore the vertical dimension. Building height influences urban ecosystems by altering wind currents and casting shadows which alter physical properties such as air temperature and evapotranspiration potential.

Structure of Urban Landscape (STURLA) spatial analysis studies have demonstrated that cities are composed of a discrete number of three-dimensional composite land classes where a small number of classes (5-10) explain the urban landscape and host distinct physical properties of the environment (Larondelle et al., 2014; Hamstead et al., 2016) and is useful for studies of urban ecosystem services (Haase et al., 2014). The urban landscape is classified by applying a fishnet grid of 120m^2^ sized pixels over a city and calculating the relative proportions of land cover elements within a pixel as derived from remotely sensed satellite imagery. STURLA classes represent combinations of the natural and built aspects of urban structure while incorporating the vertical dimension (lowrise, midrise, and highrise buildings) which gives it an explanatory advantage over most land-cover datasets. For example, class *tgplm* is a STURLA class heterogeneously composed of trees (*t*), grass (*grass*), pavement (*p*), low-rise (*l*), and mid-rise buildings (*m*). This framework allows for scalable and reproducible tracing of urban spatio-temporal patterns that can be applied across cities to identify structure function relationships. Likewise, it could easily be incorporated into models that forecast near-term changes (Dietze et al., 2018) in urban ecosystem services including and their resilience to change (Folke et al., 2002; Gunderson et al., 2012).

Urban areas host diverse microbiomes with unique selective pressures (e.g. pollution and chronic disturbance) that vary along spatial gradients (Reese et al., 2016; Mhuireach et al., 2019a); however, relationships between urban landscape heterogeneity and atmospheric microbes remain largely unexplored. Aside from outdoor air, public transit systems also host unique microbiomes. These systems are taxonomically diverse (Dybwad et al., 2012; Leung et al., 2014; Triadó-Margarit et al., 2017; Gohli et al., 2019) and represent unique ecological niches due to their high usage rate by humans and rapid turnover in occupancy. Likewise, studies of outdoor atmospheric microbiomes have until now been mostly stationary (Bowers et al., 2009; Lee et al., 2009; Mazar et al., 2016; Cáliz et al., 2018; Stewart et al., 2020) and miss the continuous biodiversity of the urban landscape. Microbiomes of both outdoor and indoor-transit air are assembled from diverse environments including soils(Bowers et al., 2009, 2011b), water (Methé et al., 1998; Linz et al., 2017), vegetation (Lymperopoulou et al., 2016), and animals (Täubel et al., 2009; Hospodsky et al., 2012; Leung et al., 2014). These communities host diverse bacterial phyla largely composed of three phyla: *Proteobacteria*, *Actinobacteria*, *Firmicutes,* and *Bacteroidetes* (Li et al., 2019; Mhuireach et al., 2019b) that vary seasonally (Cáliz et al., 2018; Du et al., 2018) and by the environment surrounding the air (Bowers et al., 2011b, 2013). Mobile monitoring (sampling while moving) of the atmosphere is increasingly used to understand patterns of particles in air (Wallace et al., 2009; Shakya et al., 2019; Deshmukh et al., 2020) and microbial diversity in a subway (Leung et al., 2014), until now it has not been applied to identifying patterns of microbial biodiversity in outdoor air relevant to the populations living in cities.

Recently, the health effects of microbial exposure in addition to pathogen identification are now being explored (Ege et al., 2011; Hanski et al., 2012; Ruokolainen et al., 2015). Low-diversity microbiomes in animals are associated with chronic-diseases including: asthma, irritable bowel syndrome (Frank et al., 2007; Round and Mazmanian, 2009), and obesity (Turnbaugh et al., 2008). Research has identified exposure to microbially diverse outdoor and built environments during childhood offers protection against asthma and immune diseases (Rook, 2009; von Hertzen et al., 2011; Mills et al., 2017). Niche occupation of the respiratory tract by diverse taxa would limit the surface area where bacteria suggested to exacerbate asthma (e.g. *Haemophilus spp.* or *Streptococcus spp.*) could settle (Ege et al., 2011). Likewise, diverse microbial exposure has been suggested to activate downstream signaling pathways of toll-like receptors that induce regulatory T-cells (Schaub et al., 2009). Given these associations, discerning patterns of diversity in urban airs holds implications for human health.

Understanding how structural elements relate to atmospheric microbial diversity and function is necessary for understanding urban ecology (Jansson, 2013), and can guide urban development in ways that utilize ecological functions to enhance environmental performance and human wellbeing. In this study we combine mobile monitoring of the atmospheric microbiome from 07/01/19 to 07/25/19 using marker gene sequencing (16s rRNA) and genome prediction to characterize the diversity and inferred functional potential in outdoor and indoor-transit air. Samples were taken outdoors by driving across 240 kilometers with eight repetitions and commuting through three train lines four times in Philadelphia PA. This is complemented with three-dimensional spatial analysis of how subtle variation in urban landscape composition influences microbial communities.

## Materials and Methods

### Site Description and Sample Collection

Philadelphia, Pennsylvania, USA is a city flanked by two bodies of water located in the Mid-Atlantic region of the United States of America with a 2018 population estimate of 1,584,138 (Lynch et al., 2014). Samples were collected on sterilized (500°C for 3 hours) quartz fiber filters using a PDR1500 Air Quality Dust Monitor (Thermo-Fisher Scientific, United States of America) where the inlet tube and mesh filter holder was cleaned with 70% ethanol before sampling. Only particles smaller than 2.5 micrometers in diameter (PM_2.5_) were collected using a particulate matter sampler with a flow rate of 1.5 liters per minute. Particulate matter was impacted on quartz fiber filters, as done in similar studies (Bowers et al., 2013; Stewart et al., 2020), where after collection the filter was removed using sterilized tweezers and stored in aluminum foil at −20 °C until DNA extraction. DNA was then stored at −80 °C until downstream analyses.

### Urban Structure and Route Generation

Urban structure was characterized in 120 m^2^ blocks using the STructure of Urban LAndscapes (STURLA) system, a multi-dimensional spatial classification system that incorporates land cover with urban building/structure height as previously described (Hamstead et al., 2016). Relative frequencies of STURLA classes were calculated and those with an abundance below 0.1% were grouped into the category “other”. STURLA across Philadelphia can be found in Figure 1. Routes were generated to include at least 50 points in each of the composite classes represented in the city STURLA classification to maximize variation in urban structure. Points were randomly selected based on a stratified random sample to ensure adequate representation of the most common composite classes. In addition, points of interest were added to the random sample including industrial sites, EPA TRI sites, and EPA air pollution monitoring station sites. Using ESRI ArcGIS 10.7.1 Network Analyst, an optimal route of approximately 480 kilometers was generated, passing through the sample points. The route was then divided into two routes—R1 and R2 for feasibility of driving within one day. The routes were loaded to a professional-grade GPS unit (Thimble Juno 3b, USA), and the van was driven along each route. Each route was replicated 4 times for a total of 8 samples; however, variation of each route exists day to day to test the role of subtle variation in the urban landscape.

**Figure 1:**
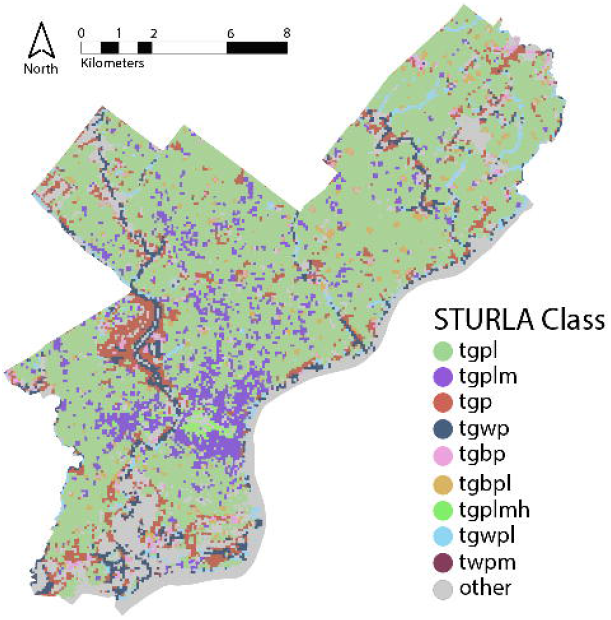
STURLA map of Philadelphia, Pennsylvania, USA where each color corresponds to a different STURLA code. T: trees, G: grass, B: bare, W: water, P: pavement, L: low-rise, M: midrise, H: highrise. The combination of these individual letters creates a STURLA code; for example, class *tgpl* is composed of various proportions of trees (T), grass (G), pavement (P), and low-rise buildings (L).

### Outdoor Sampling

Outdoor air samples (n=8) were collected using the same instrument (PDR1500) strapped to the roof of a van (~1.5 meters in height) with the inlet facing out to the side of the roof. Samples were collected from two routes (R1: Route 1, R2: Route 2) with R1 being more diverse in urban structure. The van was driven at a variety of speeds ranging from 0 km/hr - ~100 km/hr (e.g. stoplight vs highway). Average sampling period for each of the routes was 601 ± 48 minutes for each of the outdoor air sample days, collecting an average of 913.3 L ± 73.4 L of air. Variation exists within routes to address the roles of both large (between route) and small (within-route) variation in structuring the atmospheric microbiome (Figure 2). The distribution of STURLA classes throughout the city can be found in Figure 3B.

**Figure 2:**
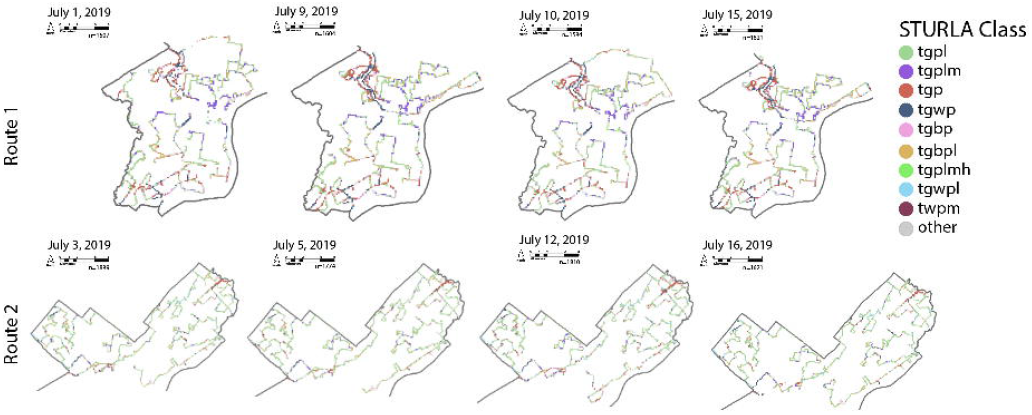
Sampling routes for each day of collection, colors correspond with STURLA class. The reader is encouraged to zoom in on each map to examine the spatial distribution of sampling and the corresponding STURLA class. T: trees, G: grass, B: bare, W: water, P: pavement, L: low-rise, M: midrise, H: highrise. The combination of these individual letters creates a STURLA code; for example, class *tgpl* is composed of various proportions of trees (T), grass (G), pavement (P), and low-rise buildings (L).

**Figure 3:**
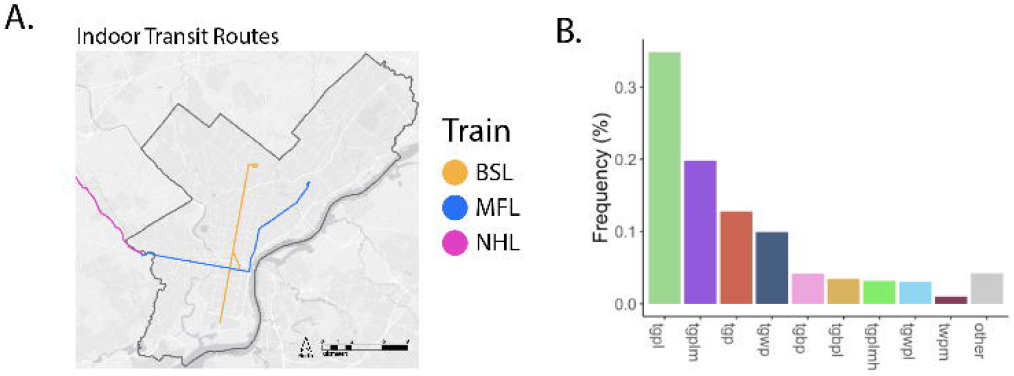
A. Routes where indoor transit system air was collected (BSL: Broad Street Line, MFL: Market-Frankford Line, NHL: Norristown Highspeed Line) B. Barplot of mean class frequency encountered while driving, colors correspond with STURLA class.

### Indoor-transit Sampling

Three indoor-transit lines (Figure 3A), the Market-Frankford Line (above/below ground), Norristown Line (above ground), and BSL (below ground) of air were collected to comprise one sample. Collection (n=4) took place inside train cars for an average 288.3 ± 21.1 minutes impacting 438.1 ± 32.4 L of air. On average ~30 people were counted in each train car during sampling.

### DNA Extraction, Sequencing, Processing, and Genome Prediction

DNA was extracted using the MoBio PowerSoil™ DNA Isolation Kit following the manufacturer’s suggested protocol. The 16S rRNA gene HV4 region for bacterial and archaeal microorganisms (515F: 5’-GTGCCAGCMGCCGCGGTAA, 806R: 5’-GGACTACHVHHHTWTCTAAT, Thermo-Fisher, Waltham, MA, USA) was then amplified by polymerase chain reaction (PCR) using the Earth Microbiome Project’s protocol (Earth Microbiome Project, 2016). A positive control of *E. coli* and a negative control of deionised water were used and no contamination was found in either. Sequencing was performed at MR DNA (www.mrdnalab.com, Shallowater, TX, USA) using the Illumina MiSeq platform. DNA sequences were analyzed and statistical analysis was conducted using QIIME 2 (Bolyen et al., 2019) following the *Moving Pictures* tutorial code and R (3.6). Briefly, raw reads were demultiplexed, denoised using the DADA2 algorithm (Callahan et al., 2016), and truncated at 180 basepairs.

Sequences were rarefied to 1,55,547 sequences aligned to the SILVA 132 99% reference taxonomy database (Quast et al., 2013) using a Naive Bayes classifier to amplicon sequence variants (ASVs). Sequences were aligned using mafft (Katoh and Standley, 2013) followed by construction of an unrooted phylogenetic tree using FastTree (Price et al., 2009). Alpha diversity metrics including observed number of ASVs, Pielou’s evenness, and Faith’s Phylogenetic Diversity (Faith, 1992) were calculated. Beta diversity was calculated using unweighted UniFrac for phylogenetic analysis (Lozupone and Knight, 2005). Unweighted UniFrac was chosen as it is qualitative and thus makes our analyses less sensitive to differences in sampling volume and allows comparability to qualitative functional inference (e.g. functional Bray-Curtis PcoA). PICRUSt 2 (Douglas et al., 2019) was used to infer functional genes using the maximum parsimony hidden-state prediction methodology (Louca and Doebeli, 2018) as done in other studies of the atmospheric microbiome (Pan et al., 2019; Stewart et al., 2020). Functional gene predictions were rarefied to the minimum number of features found between all samples. Beta diversity for functional gene predictions was conducted using Bray-Curtis dissimilarity and Principal Coordinates Analysis (PcoA).

### Data Analysis

Identification of potentially pathogenic taxa was conducted comparing the average 40 most relatively abundant ASVs with a reference database of bacterial strains that are known human pathogens (UK Health and Safety Executive Advisory Committee on Dangerous Pathogens). Taxa representing less than 0.01 % of the pathogenic subset of taxa were grouped into the category “Other”. Likely source environment for the top 40 ASVs was determined through a literature review (Stewart et al., 2020) (supplemental references). A taxon found in more than one environment was categorized as a “generalist” as done in previous research (Stewart et al., 2020). For this the relative abundance of each taxon was coded based on their likely source environment. Coded environment was then used for statistical analysis as seen below.

Bray-Curtis dissimilarity for functional gene predictions, source environment, and potentially pathogenic taxa was calculated. Kruskal-Wallis tests were used to test for significant differences indoor-transit, outdoor air, and variation between R1 and R2 in diversity. PERMANOVA tests identified if community structure (phylogenetic, by source environment, predicted functional genes, and potentially pathogenic taxa) varied between air types and route driven. Taxon reoccurrence was measured by taking the sum of the presence of a taxon in outdoor and indoor-transit samples and breaking them into equal thirds ranked by number of occurrences across all samples.

Shannon diversity (H) was used to calculate the diversity of STURLA classifications encountered (Shannon and Weaver, 1963). Richness of land cover was measured by taking the sum of the number of unique STURLA classes passed through on each route. These two metrics are used as proxies for how many STURLA classes (e.g. as one walks throughout a city) and which classes in a city are to be associated with microbial diversity. Dissimilarity in routes was calculated using the Bray-Curtis dissimilarity metric (Beals, 1984). Permutational correlations (to account for spatial and temporal autocorrelation) with Bonferroni correction were used to test for relationships between STURLA diversity and microbial structure/function using relative proportions of STURLA classes (as this accounts for differences in total driving time). A mantel test was employed to test for correlation between microbial beta diversity and STURLA dissimilarity.

### Data availability

DNA sequences have been deposited to the National Institute of Health Sequence Read Archive under accession number: SRR11612565. Code can be found at https://github.com/thecrobe/STURLAMicrobiome.

## Results

### Microbial Community Structure and Function

Molecular analysis from mobile monitoring revealed diverse communities of 337 amplicon sequence variants (ASVs, a proxy for species). 22 phyla were observed that varied significantly in community structure (Figure 4A, *p*=0.001) and inferred functional potential (Figure 4B *p=*0.002) between outdoor and indoor-transit air as well as between outdoor route driven. Outdoor communities were seen to host reoccurring taxa less frequently than indoors (Figure S1). A diverse set of metabolic genes were identified including carbon fixation, sulfate reduction/assimilation/oxidation, nitrate reduction, and aromatic compound degradation. No virulence genes were observed. Despite differences in composition and inferred functional potential, both indoor-transit and outside air hosted reoccurring taxa similarly (*p=*0.9631, Figure S1) where the majority of ASVs in outdoor and indoor-transit were found in all samples. No phylum was unique to a single sample, air type, or route (Figure 4C) thus suggesting that the atmospheric microbiome in general is panmictic; however may vary between indoor and outdoor airs.

**Figure 4 :**
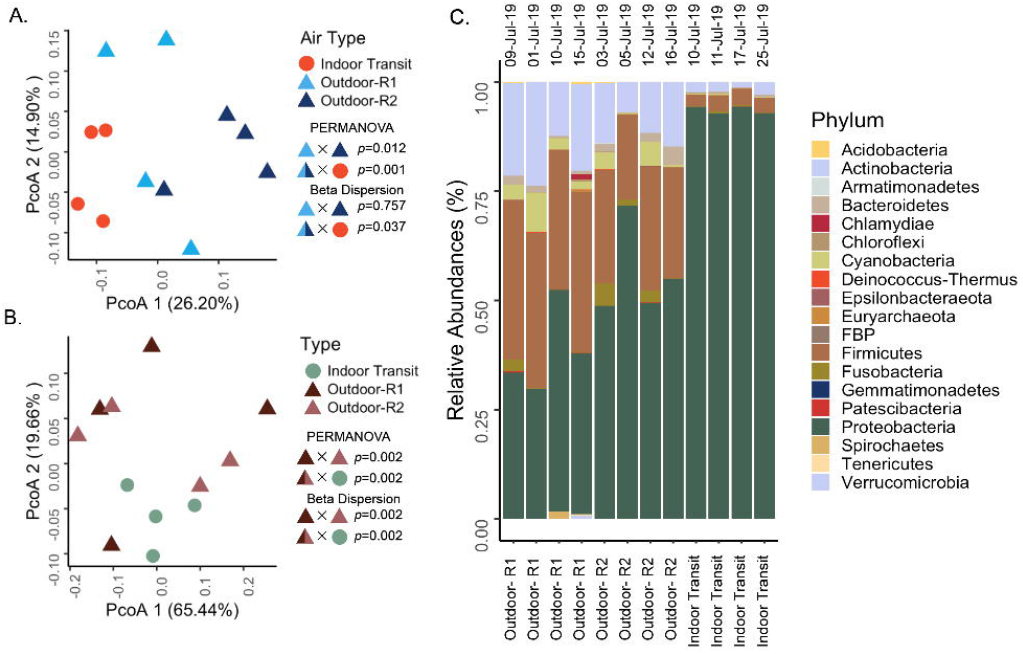
A. Principal Coordinates Analysis (PcoA) of phylogenetic beta-diversity (between sites) using unweighted UnifFrac distances with air type colored by site and shaped by indoor (circle) and outdoor (triangle) air. Shape/Color X Shape/Color indicates the factors used in hypothesis testing with the test above and p-value to the right. B. Principal Coordinates Analysis (PcoA) of inferred functional potential beta-diversity (between sites) using Bray-Curtis dissimilarity with air type colored by site (green: indoor transit, dark brown: outdoor route 1, light brown: outdoor route 2) and shaped by indoor (circle) and outdoor (triangle) air. Shape/Color X Shape/Color indicates the factors used in hypothesis testing with the test above and p-value to the right. It should be noted the split color is for visualization and statistical tests were conducted on outdoor ~ indoor samples. C. Relative abundances of phyla identified in samples separated by air type (outdoor route 1 vs outdoor route 2 vs indoor transit) with collection date above each column.

Indoor air hosted almost double the number of rare taxa (49 ASVs), an ASV found in just one sample, than outdoor air (26 ASVs). Outdoor air hosted significantly more phylogenetic (*p=*0.006, Figure 5A) and functionally diverse (*p=*0.002, Figure 5B) communities as compared to indoor-transit air. Furthermore, outdoor communities were more rich (*p=*0.045, Figure 5C), and evenly distributed (p=0.007, Figure 5D). The 40 most abundant ASVs accounted for an average 86.01% (coefficient of variance: 11.06%) of all taxa across all samples were identified as likely originating from animals, soil, or were generalists (Figure 6A). Enterobacteriaceae (likely from mammalian gastro-intestinal systems) were on average the most abundant animal-associated microorganisms and were more variable outdoors (16.71% ± 15.01%) than indoors (90.5% ± 1.7%).

**Figure 5.**
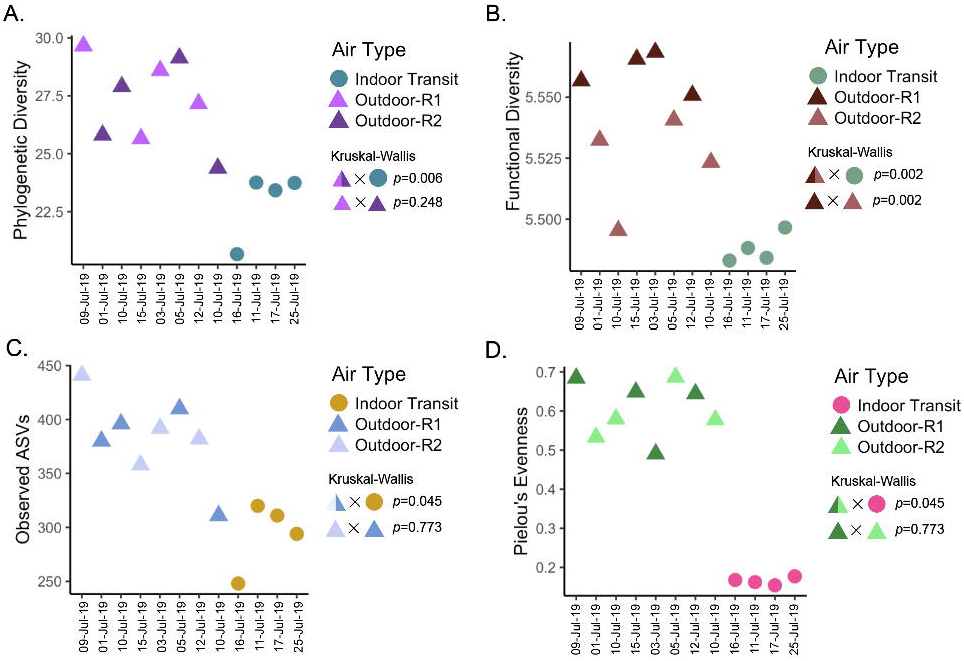
Alpha Diversity Metrics separated by air type (circle: indoor transit, triangle: outdoor) and route driven for outdoor samples (color) with date of collection on the x axis. Shape/Color X Shape/Color indicates the factors used in hypothesis testing with the test above and p-value to the right. It should be noted the split color is for visualization and statistical tests were conducted on outdoor ~ indoor samples. A. Faith’s phylogenetic diversity (PD). B. Inferred functional diversity calculated using the Shannon index on PICRUSt functional pathways. C. Observed number of ASVs. D. Pielou’s evenness metric.

**Figure 6.**
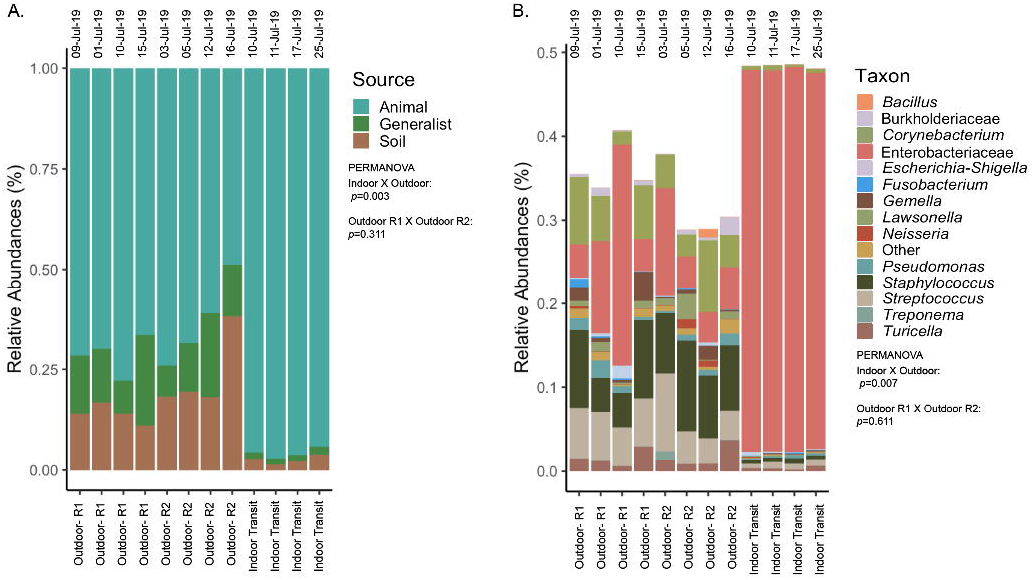
A. Relative abundances of sources identified in samples separated by air type on the bottom (outdoor route 1 vs outdoor route 2 vs indoor transit) with collection date above each column. Hypothesis test results and factors are under the legend. B. Relative abundances of potentially pathogenic taxa identified in samples separated by air type on the bottom (outdoor route 1 vs outdoor route 2 vs indoor transit) with collection date above each column. Taxa in abundance smaller than 0.01% were grouped in the category “other” for visualization only. Hypothesis test results and factors are under the legend.

While indoor transit air was dominated by Enterobacteriaceae, the skin associated genus *Staphylococcus* (mean: 14.22%, coefficient of variance: 30.62%) made notable contributions to the potentially pathogenic fraction of outdoor air communities and was more variable in abundance across samples as compared to indoors (mean: 0.01%, coefficient of variance: 18.90%). Plant pathogen containing genus *Ralstonia* represented the most abundant taxa sourced to soil and were in greater abundance and less variable outdoors (mean: 17.56%, coefficient of variance: 9.48%) than indoors (mean: 2.49%, coefficient of variance 39.15%). Lastly, the most abundant generalist ASVs were identified as the Oxyphotobacteria, the only group of bacteria that can carry out oxygenic photosynthesis, were also more abundant and variable outdoors (mean: 4.16%, coefficient of variance: 82.73% than indoors (mean: 0.24%, coefficient of variance: 70.87%). Taxonomic composition by source environment significantly differed from outdoor and indoor-transit air (*p=*0.003) but not by route driven (*p=*0.311) Proportions of source environments were not uniform between outdoor and indoor-transit air as the most abundant ASVs in outdoor air were attributed to more evenly distributed source environments.

A total of 30 Amplicon Sequence Variants (ASVs) were identified as being potentially pathogenic where, on average, these taxa composed (mean: 33.89%, coefficient of variance: 19.0%) of outdoor air and (mean: 48.41%, coefficient of variance 21.1%) of indoor-transit microbial communities (Figure 6B). The majority of these taxa are commensal human bacteria that are also opportunistic pathogens. *Corynebacterium spp.,* and *Pseudomonas spp.,* were absent in indoor-transit air. Structure and taxonomic composition of potentially pathogenic taxa was significantly different between indoor-transit and outdoor air (*p=*0.007) but not between route driven outdoors (*p=*0.661).

### The Three-Dimensional Landscape and Biodiversity

Classification of the urban landscape identified 42 STURLA classes encountered while driving with *tgpl (trees, grass, pavement, lowrise buildings)* being most frequent (Figure 2, larger maps found in Supplemental Figures 5-12).The majority of the urban landscape from routes driven was homogenous in class composition where 92.16 % ± % 1.43 % was explained by 9 STURLA classes (Figure 3B). Despite this homogeneity, routes significantly differed (*p=*0.0364) in STURLA classes encountered each day. Diversity of STURLA classes encountered was significantly correlated with phylogenetic diversity (Rho: 0.74, *p=* 0.045) and observed number of ASVs (Rho: 0.83, *p=* 0.011) but not with community evenness (Rho: 0.21, *p=* 0.620). We found that variation of the STURLA classes was significantly associated with beta diversity of microbial communities (Mantel Test, r: 0.341, *p=*0.04) At the individual STURLA class level, inferred functional diversity was significantly correlated with *tgpl*, *tgplmh*, *other* ( all *p* <0.05) and *tgp, tgwp, tgbp,* (all *p* < 0.1) (Figure 7) as was the number of observed ASVs and phylogenetic diversity with *tgpl* and *tgplmh*. Furthermore, diversity of STURLA classes by day was strongly, albeit insignificantly, correlated with 14 metabolic pathways (Supplemental Figure 2), 25 biosynthesis pathways (Supplemental Figure 3), and 10 degradation pathways (Supplemental figure 4). Class richness was also correlated with 3 metabolic pathways, 9 biosynthesis pathways, and 4 degradation pathways.

**Figure 7:**
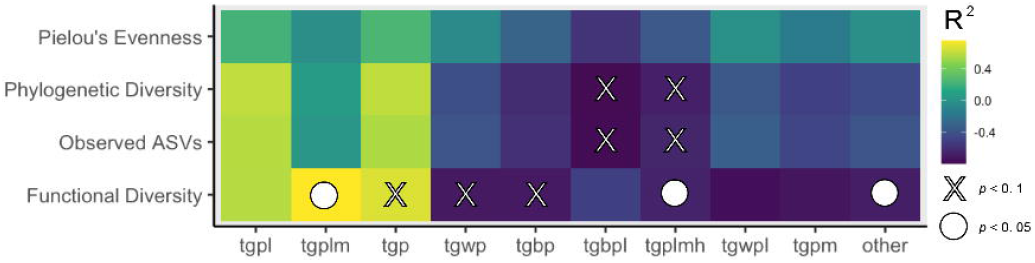
A. Heatmap of Permutational correlation coefficients where lighter colors indicate magnitude of the correlation. X indicates a statistically significant p-value after Bonferroni correction. STURLA Code elements: t: trees, g: grass, p: pavement, b: bare, w: water, l: lowrise, m: midrise, h, highrise.

## Discussion

The high level of diversity observed in outdoor particulate matter is likely attributed to the unrestricted dispersal by wind at both the local (within-city) (Pankhurst et al., 2012) and intercontinental scales (Smith et al., 2013). This diversity may also be a function of high temporal variation at short time scales (i.e. days) of microbial biodiversity previously found in the atmospheric microbiome (Stewart et al., 2020) and should be considered when interpreting these results. This is attributed to sampling across diverse landscapes that included water features, greenspace, unpaved land as well as a dense urban core consisting of the built environment where humans and animal companions primarily walk and drive outside. For atmospheric communities it is expected that dispersal is the dominating process affecting community assembly as opposed to diversification, selection, and drift (Vellend, 2010; Nemergut et al., 2013). We also found a relatively small number of ASVs as compared to other environments (e.g. soil); however, this is consistent with estimates of atmospheric microbial biodiversity that has found 29-690 phylotypes pers sample (Bowers et al., 2013). Our finding that communities host reoccurring taxa over time in both outdoor and indoor-transit air suggests that atmospheric microorganisms are not endemic and have cosmopolitan distributions in their environment, likely a function of the large populations of microorganisms combined with their small size (Wilkinson et al., 2012). Despite this, the significant differences in community composition by air type also suggests that the mixing is dispersal limited (e.g. greater mixing outside as opposed to a largely enclosed built environment). When outdoors, aerosolized microbes are blocked by buildings, trees, and everything in the urban landscape; however they have a wider possible geographic range (a function of wind dispersal) than indoors. For example, a microbe aerosolized from a plant leaf can move in all directions, including vertically where it could be lifted and transported long distances (Smith et al., 2013). Being able to across wide geographic ranges allows endemic microbes to escape their native geographic ranges. While inside a building such as a subway or train station wind speeds do not reach levels found outdoors. Likewise, indoor microbes have an obstacle, exits/entrances. The microorganisms would have to exit the system before it could enter outdoor air.

Furthermore, in semi-closed environments such as indoor-transit systems, physical dispersal limitation (e.g. a pillar or turnstile) should increase residence time for microbes (Adams et al., 2013; Kembel et al., 2014) and influence microbial diversity. The closed environment (aside from exits) of the indoor-transit system combined with high human occupancy seems to enrich the taxa brought into the system by its riders as it does in other structures of the built environment (Meadow et al., 2014). It is expected to find bacteria disperse into and throughout the indoor-transit environment by three primary mechanisms: (1) contact with indoor surfaces; (2) particle emission from breath, clothes, skin and hair; and (3) resuspension of dust containing other microbial-loaded particles (Meadow et al., 2015). Given that humans carry 10^12^ microorganisms on the epidermis and 10^14^ microorganisms in the alimentary tract (Luckey, 1972) and shed ~ 10^6^ particles (>0.5 μm in diameter) an hour (You et al., 2013; Bhangar et al., 2016) we likely serve as the dominant source of microorganisms in indoor-transit air.

Complementing our findings of higher proportions of animal-associated taxa in indoor-transit air, it has been seen that human occupancy increases the load of animal-associated bacteria in the built environment. This includes subways (Leung et al., 2014; Hsu et al., 2016) university classrooms (Hospodsky et al., 2012; Meadow et al., 2014), public restrooms (Flores et al., 2011), dust (Täubel et al., 2009), and kitchens (Flores et al., 2013). Outdoor air microbes also seem to enter into the transit system, albeit to a lesser extent, given there were no unique phyla identified outside that were not encountered in transit air. Furthermore, architectural design (Kembel et al., 2012, 2014) has been seen to influence microbial community structure and acts as a physical barrier differentiating outdoor versus indoor-transit air which may account for observed variation in diversity in this study.

High human occupancy in small spaces such as train cars combined with our finding of almost half of the microbiome being potentially pathogenic may alter the health state of the lung and respiratory tract microbiome. Despite the lack of specific inferred virulence genes, shifts in microbial composition can have negative health effects as an emergent property. A decrease in microbial diversity, richness and evenness, as we found in indoor-transit air compared to outdoor air, are frequent features of chronic lung inflammation (Hilty et al., 2010; You et al., 2013). The presence of microbial derived molecules influences immune systems through Toll-like receptor recognition even when a microbe is dead. These molecules include bacterial lipopolysaccharide (Faure et al., 2000; Ibeagha-Awemu et al., 2008), flagellin (Hayashi et al., 2001), and lipoproteins(Takeuchi et al., 2001). Thus, dead microbes that lack pathogenicity genes may still have negative health outcomes upon inhalation by activating the immune system. Previous research has demonstrated that approximately 80% of particulate matter (PM_2.5_) in subway systems deposits in human respiratory tract and biologically stresses the immune system and cultured lung cells (Karlsson et al., 2005; Bachoual et al., 2007). This combined with the high percentages of potentially pathogenic taxa in indoor-transit air may have negative health effects on those with unstable lung microbial communities such as individuals who have a recurring cough, mechanically disrupting microbes, or have decreased diversity due to antibiotic use (Klepac-Ceraj et al., 2010; Zhao et al., 2012). In contrast, the lower proportions of potentially pathogenic taxa and relatively higher diversity of outdoor air (Ege et al., 2011; Lynch et al., 2014) has been linked to decreased rates of asthma. The influence of outdoor air may be less than that of the indoor-transit system as 90% of urban individuals spend their time indoors (Klepeis et al., 2001).

One possible benefit of spending time outdoors aside from aesthetics and recreation (Soga and Gaston, 2016) may be improving respiratory health. It should be noted that these are suggestive findings that would benefit from empirical measurements from qPCR or other functional analyses. Effects of microbial presence in the atmosphere on respiratory tract colonization is currently unknown; however, it is suggested to be largely determined by processes including birth, death, immigration, and emigration (Whiteson et al., 2014; Willis et al., 2020). Recent studies have also observed relationships between post-mortem microbial biodiversity on the human body (e.g. nose, rectum, and eyes) that suggest urban inhabitants are influenced by their local atmospheric microbiome (Pearson et al., 2019, 2020) and population demographics (Pechal et al., 2018). Likewise, black carbon (Janssen et al., 2011), particulate matter (Dockery et al., 1993), and other pollutants may play a role in counteracting the benefits of microbial exposure.

While it is known that urban and rural/agricultural air host distinct microbial communities (Shaffer and Lighthart, 1997; Bowers et al., 2011a; Leung et al., 2014; Stewart et al., 2020) the role of within city variation where there are less obvious contrasts in landscape structure is largely unstudied.

Likewise, understanding the biogeography of microbial communities in cities offers a way to investigate colonization of urban residents by microbes. Recent studies investigating these relationships have shown that microbial diversity of deceased individuals is significantly and positively correlated with greenness (Pearson et al., 2019, 2020). While this study does not investigate individual human microbiomes, we do find the opposite relationships with classes that host greenness, specifically *tgbpl* and *tgplmh.* Both of these STURLA classes host greenness (*t*) and the interaction between greenspace and other urban elements (*l*, *m,* and *h*) suggest that relationships between urban structure and atmospheric microbial biodiversity are an emergent property of the built and natural environment. Likewise as trees across the majority of Philadelphia’s urban landscape (*tgpl*) measuring greenness alone does not account for the true variation in the 3D landscape.

More associations between biodiversity and urban structure were seen with STURLA diversity as opposed to STURLA richness, suggesting the composition of heterogeneous 3D landscapes has greater influence on microbiome composition than the number of times the same landscape is encountered throughout a day. Likewise more correlations with urban structure were observed with inferred function genes which suggests that STURLA classes may host similar taxa phylogenetically and taxonomically but be tailored to specific functions. This may be a result of environmental filtering of microbial communities on the land (Lauber et al., 2013) below the atmosphere that are then aerosolized. Further investigations of microbial biodiversity in the atmosphere above and on the land within STURLA classes could provide urban planners with buildable units of land to increase functional diversity (Allison and Martiny, 2008) and urban resilience (Gómez-Baggethun and Barton, 2013; Jansson, 2013; McPhearson et al., 2016). For example, significant positive correlations between inferred functional diversity and STURLA class *tgplm* were observed while a negative relationship was found with *tgplmh*. This research provides further evidence that biodiversity is influenced by urban structure composed of the built and natural environment (Pearson et al., 2019). It should be noted that the sampling was heterogenous as samples were collected while stationary (e.g. stoplight) as well as when moving (e.g. city streets and highways) as well as the density of cars and people on the streets. These findings suggest that different structures of the urban landscape host taxonomically and functionally distinct microbiomes in their aerosols.

In a rapidly urbanizing global community, humans spend the majority of their time in urban landscapes of varying landscape structure, density and configuration. While both outdoors and in transit systems, we constantly emit and interact with microbes sourced from diverse environments that may influence our well-being. In this study we found that microbial communities vary between outdoor and indoor-transit air with the former being more diverse and associated with the structure of the urban landscape. Likewise indoor-transit systems hosted largely homogenous communities with higher proportions of potentially pathogenic taxa likely emitted from animals. Limitations include not analyzing the fungal and viral fractions of atmospheric particles, use of inferred functional pathways instead of directly measuring gene expression or metagenomes (Steele and Streit, 2005), and testing the PM_2.5_ subset of the atmospheric particles as opposed to total suspended particles (TSP). Likewise, sampling efficiency/length/speed, local pollutant conditions inhibiting molecular analyses and seasonality may be latent factors influencing our results. Future studies may consider exploring the variations of relationships to microbial diversity within a STURLA class as test the influence of within-season and between-season variation.

## Acknowledgements

This research was supported by NSF Grant 1832407. This manuscript has been released as a preprint at Biorxiv under DOI https://doi.org/10.1101/2020.06.17.157651 (Stewart, 2020)

We appreciate the help from Gerardo Bandera and Jennifer Santoro for analyses and Radley Reist for sample collection. Collina Strada’s sustainable fashion line is credited with colors used in graphics. We thank the editor and two reviewers for their time and energy.

## Conflict of Interest

The authors declare no conflict of interest.

## Notes

https://github.com/thecrobe/STURLAMicrobiome

### Competing Interest Statement

The authors have declared no competing interest.

