## Supplemental for "Outdoor Atmospheric Microbial Diversity is Associated with Urban Landscape Structure and Differs from Indoor-Transit Systems as Revealed by Mobile Monitoring and Three-Dimensional Spatial Analysis"

800 Lancaster Avenue, Villanova, PA 19085, USA

Department of Geography & the Environment

Villanova University

^a^Department of Geography and the Environment, Villanova University, Pennsylvania, USA

^b^Department of Ecological Science, Vrije Universiteit Amsterdam, Amsterdam, The Netherlands

Supplemental:

Supplemental Table 1: Sample information with ID, air type, date, total sampling time in minutes, and volume of air collected.

| *Sample ID* | *Date* | *Air Type* | *Total Time (Minutes)* | *Volume*  *(Liters)* |
| --- | --- | --- | --- | --- |
| *P1* | 1-Jul-19 | Outdoor-R1 | 635 | 965.2 |
| *P2* | 3-Jul-19 | Outdoor-R2 | 655 | 995.6 |
| *P3* | 5-Jul-19 | Outdoor-R2 | 536 | 814.72 |
| *P4* | 9-Jul-19 | Outdoor-R1 | 645 | 980.4 |
| *P5* | 10-Jul-19 | Outdoor-R1 | 632 | 960.64 |
| *P6* | 12-Jul-19 | Outdoor-R2 | 604 | 918.08 |
| *P7* | 15-Jul-19 | Outdoor-R1 | 562 | 854.24 |
| *P8* | 16-Jul-19 | Outdoor-R2 | 538 | 817.76 |
| *S1* | 10-Jul-19 | Subway | 298 | 452.96 |
| *S2* | 11-Jul-19 | Subway | 279 | 424.08 |
| *S3* | 17-Jul-19 | Subway | 264 | 401.28 |
| *S4* | 25-Jul-19 | Subway | 312 | 474.24 |

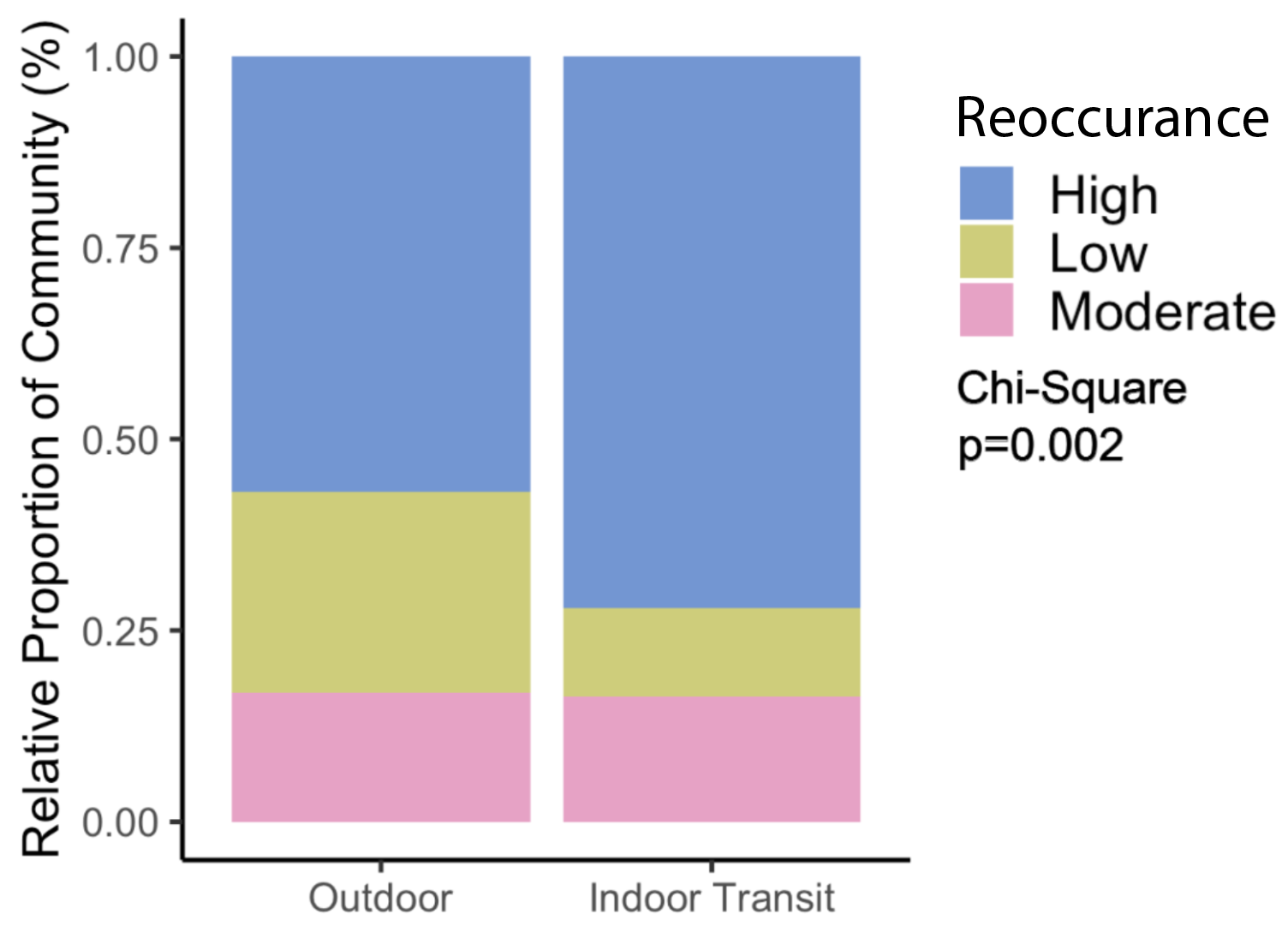

Supplemental Figure 1: Dispersal distributions of ASVs between all samples separated by air type. Colors denote degree of dispersal as measured by the sum of ASVs found in the top, middle, and bottom 33% of the community. A Chi-Square test was used to see if dispersal patterns differ by air type (p=0.002).

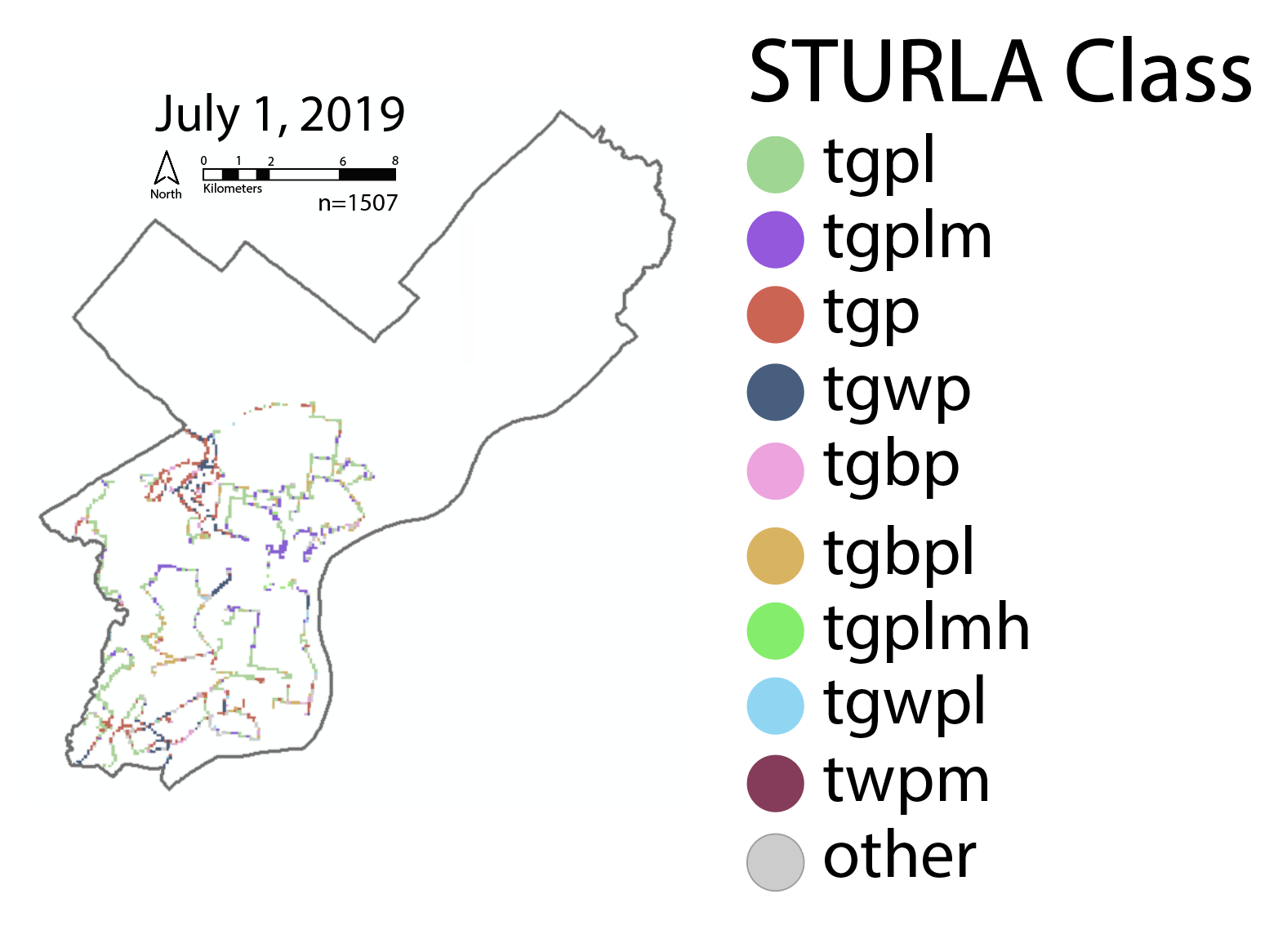

Supplemental Figure 2: Map of sampling route for July 1, 2019 where color indicated the STURLA code. T: trees, G: Grass: W: water, B: bare, P: Pavement, L: low-rise, M: midrise, H: highrise.

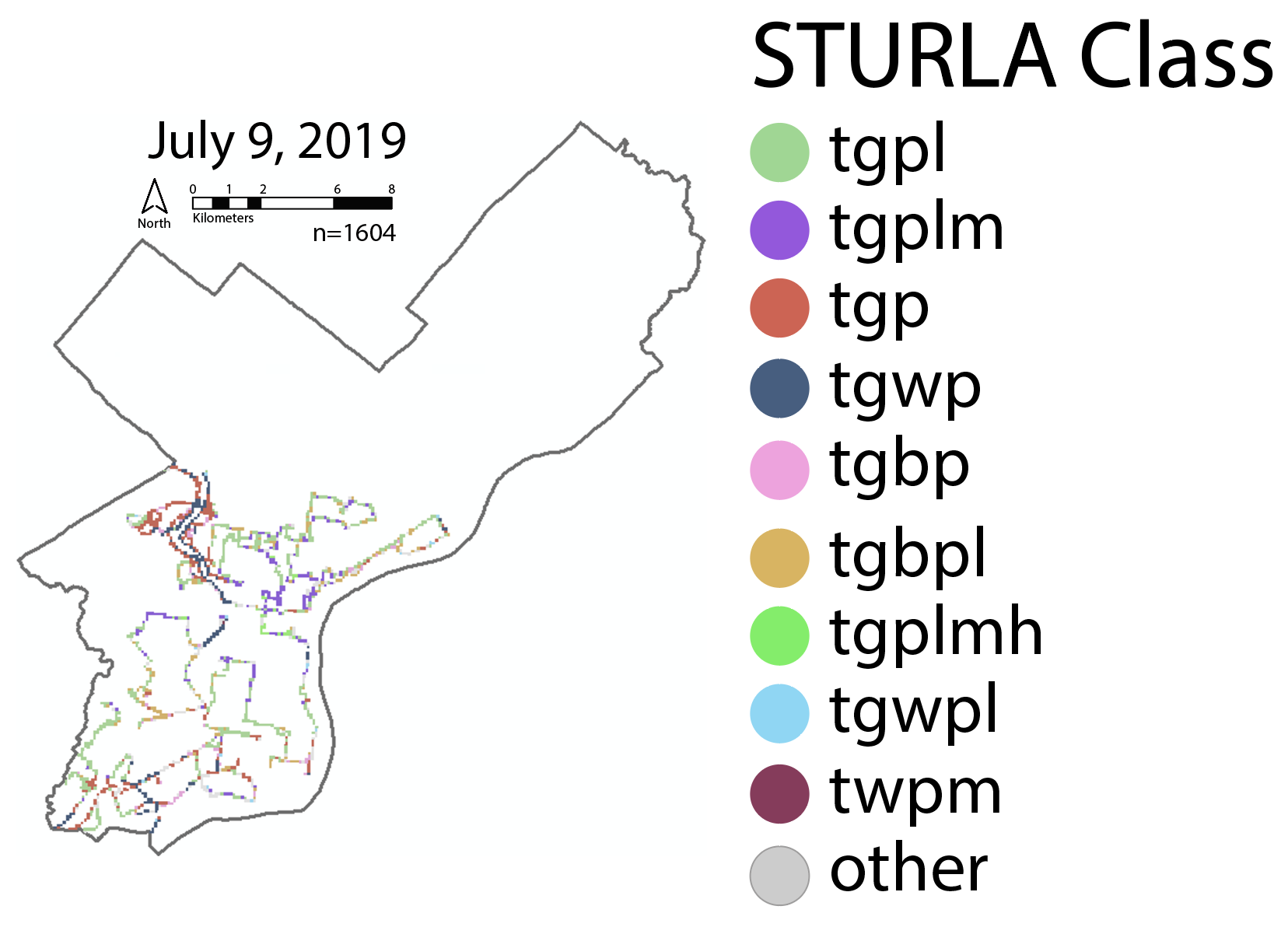

Supplemental Figure 3: Map of sampling route for July 9, 2019 where color indicated the STURLA code. T: trees, G: Grass: W: water, B: bare, P: Pavement, L: low-rise, M: midrise, H: highrise.

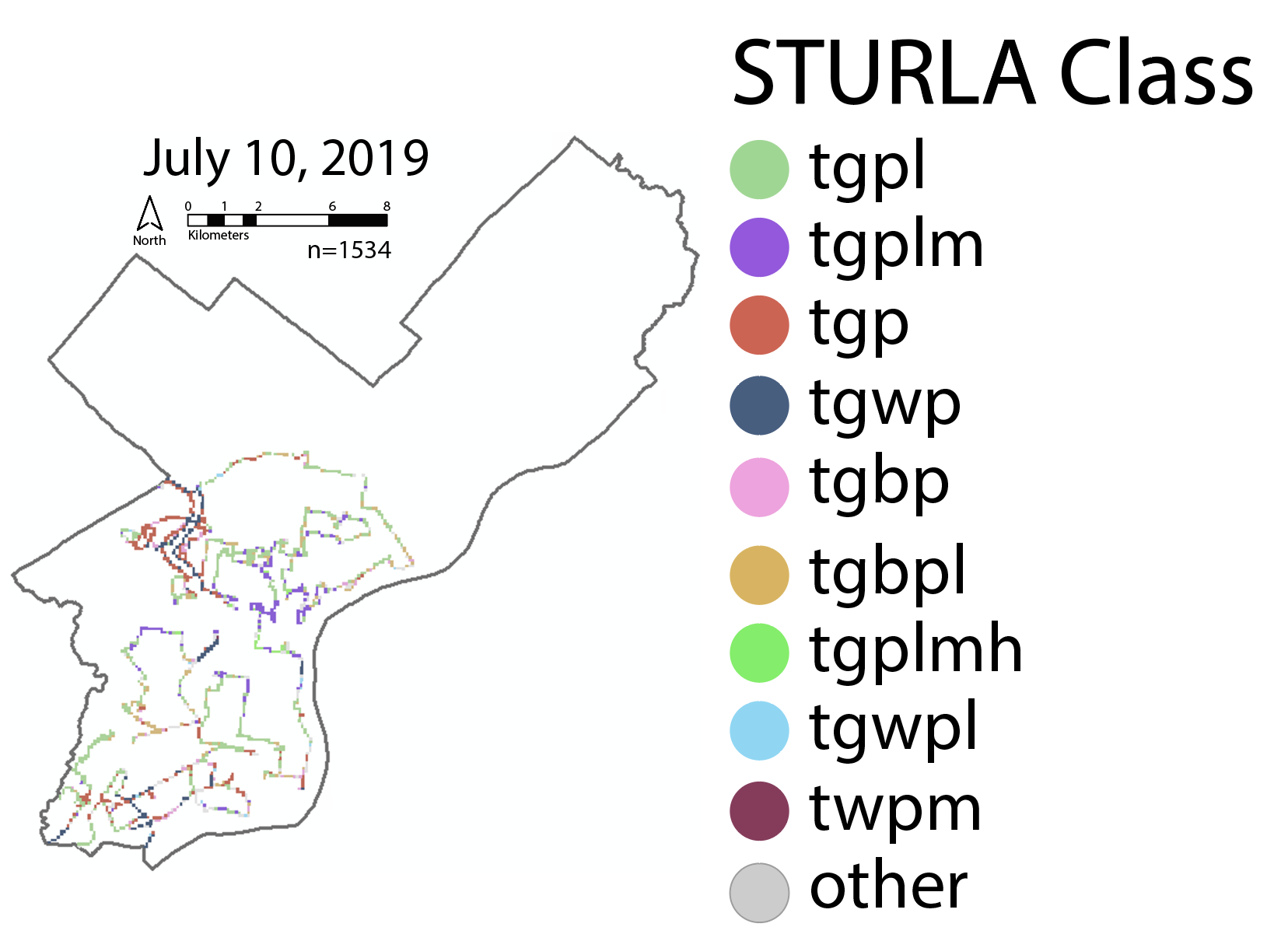

Supplemental Figure 4: Map of sampling route for July 10, 2019 where color indicated the STURLA code. T: trees, G: Grass: W: water, B: bare, P: Pavement, L: low-rise, M: midrise, H: highrise.

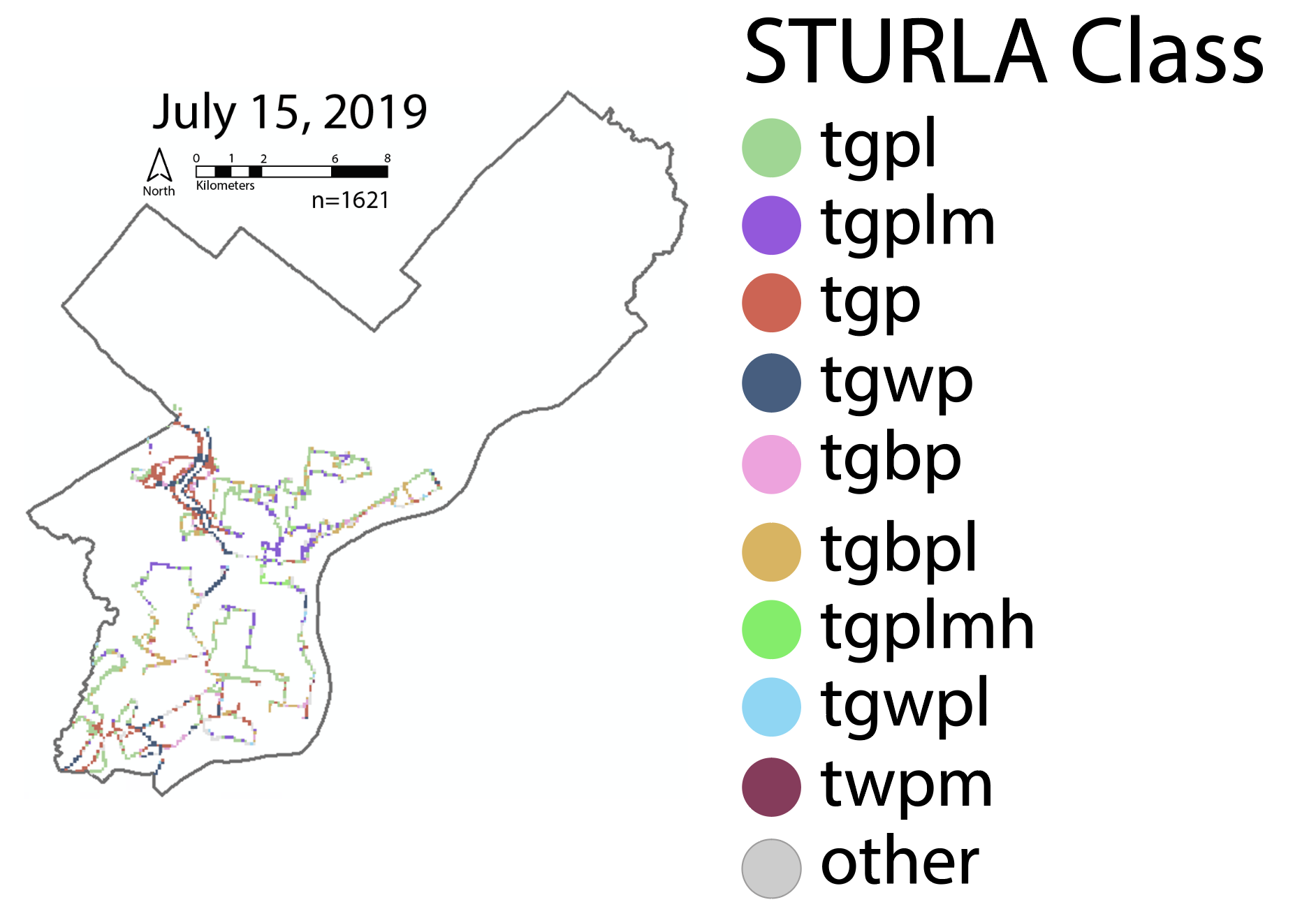

Supplemental Figure 5: Map of sampling route for July 15, 2019 where color indicated the STURLA code. T: trees, G: Grass: W: water, B: bare, P: Pavement, L: low-rise, M: midrise, H: highrise.

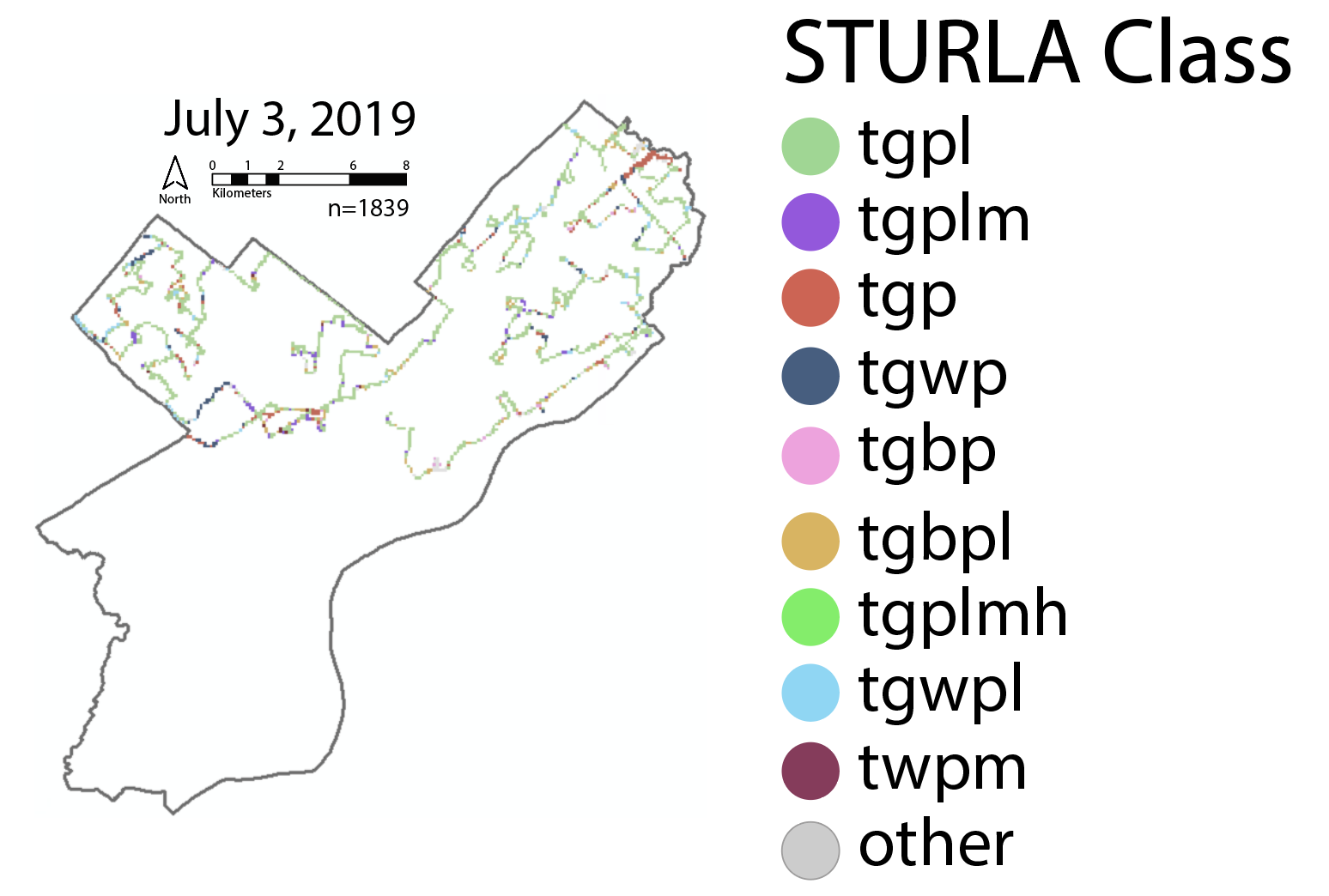

Supplemental Figure 6: Map of sampling route for July 3, 2019 where color indicated the STURLA code. T: trees, G: Grass: W: water, B: bare, P: Pavement, L: low-rise, M: midrise, H: highrise.

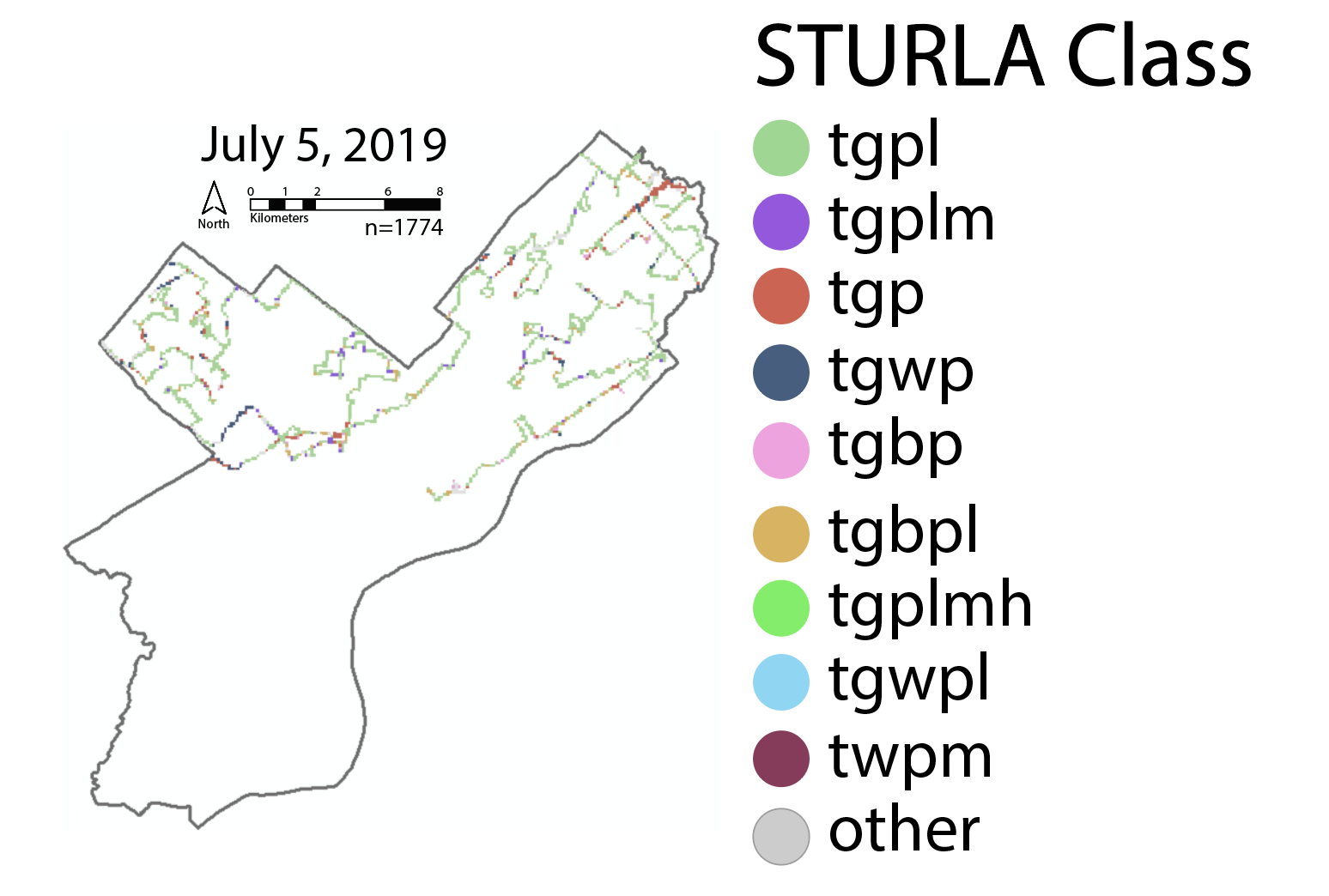

Supplemental Figure 7: Map of sampling route for July 5, 2019 where color indicated the STURLA code. T: trees, G: Grass: W: water, B: bare, P: Pavement, L: low-rise, M: midrise, H: highrise.

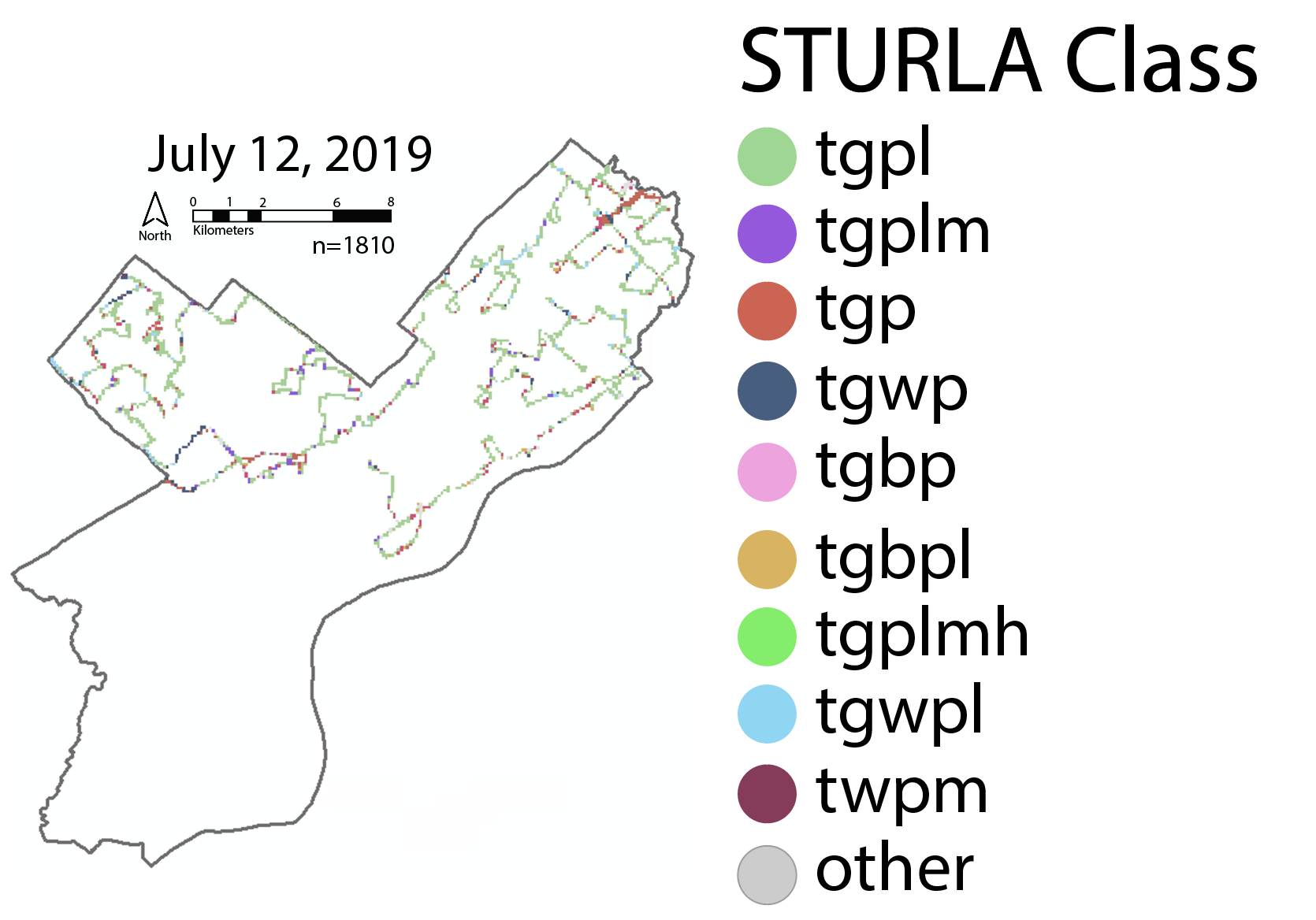

Supplemental Figure 8: Map of sampling route for July 12, 2019 where color indicated the STURLA code. T: trees, G: Grass: W: water, B: bare, P: Pavement, L: low-rise, M: midrise, H: highrise.

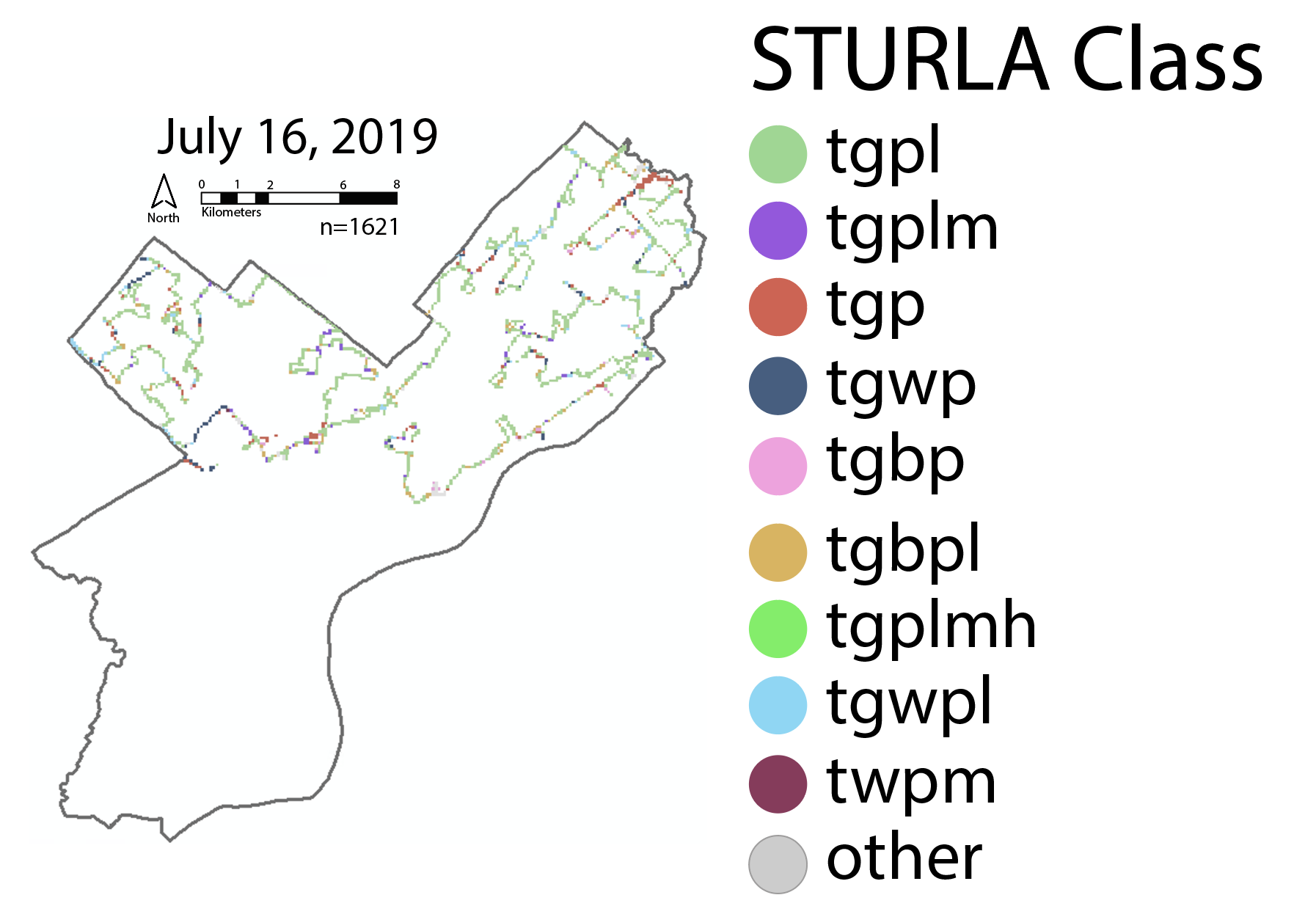

Supplemental Figure 9: Map of sampling route for July 16, 2019 where color indicated the STURLA code. T: trees, G: Grass: W: water, B: bare, P: Pavement, L: low-rise, M: midrise, H: highrise.

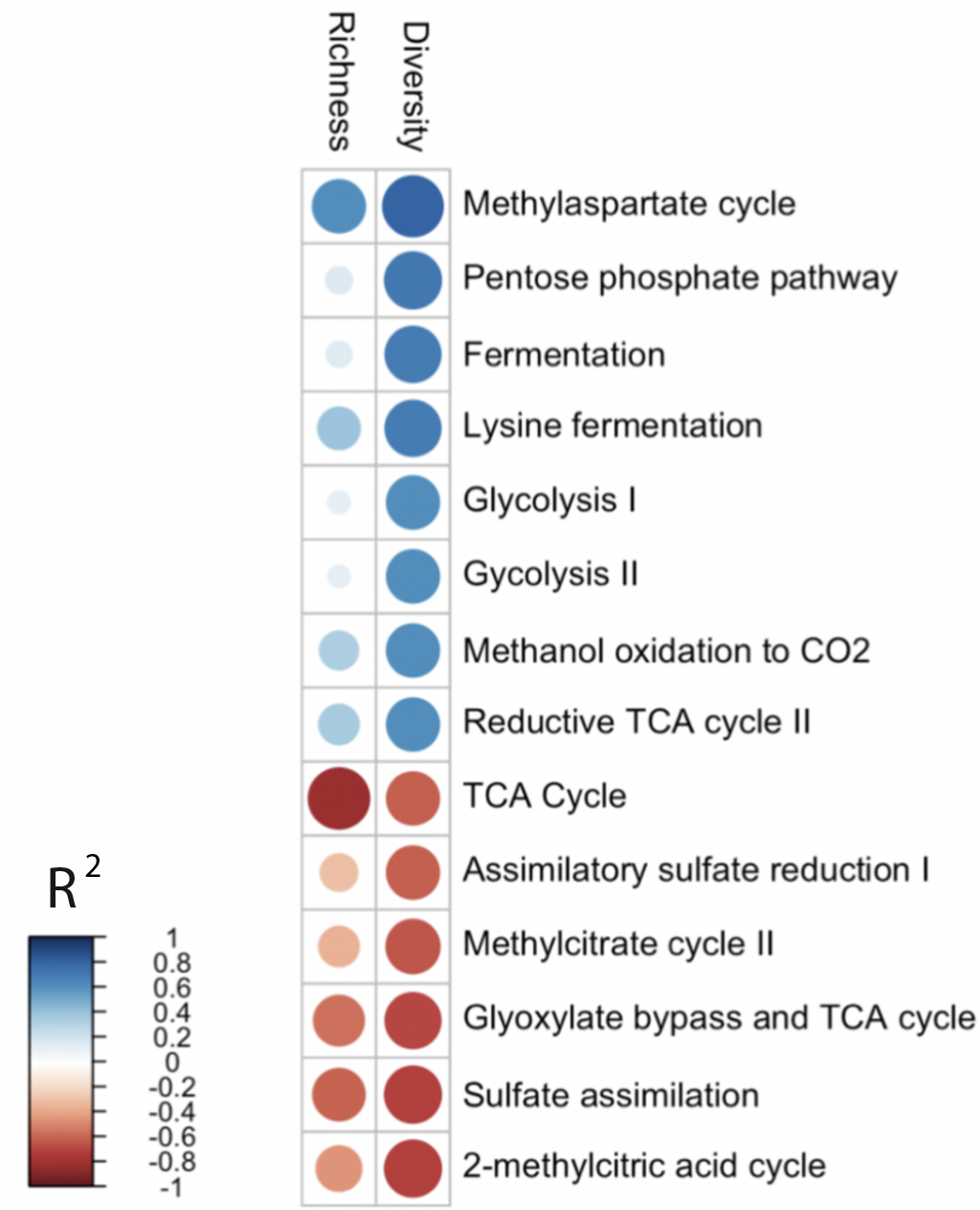

Supplemental Figure 2. Permutational correlations between metabolism associated functional genes identified using PICRUST2 and diversity and richness of STURLA classes. Strong correlations are set at a threshold of |0.70| with Bonferroni corrections (no significant relationships found).

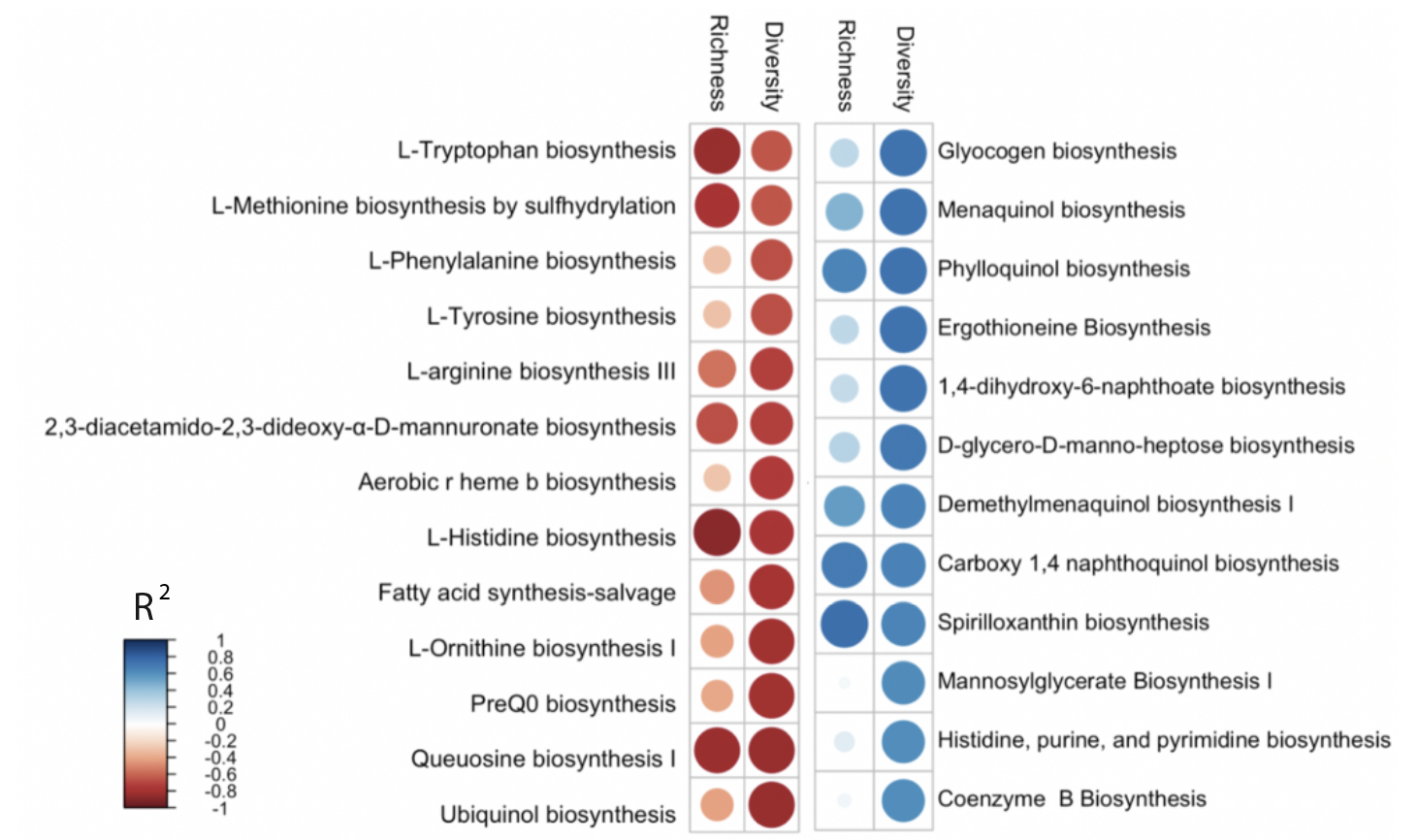

Supplemental Figure 3. Permutational correlations between biosynthesis functional genes identified using PICRUST2 and diversity and richness of STURLA classes. Strong correlations are set at a threshold of |0.70| with Bonferroni corrections (no significant relationships found).

 
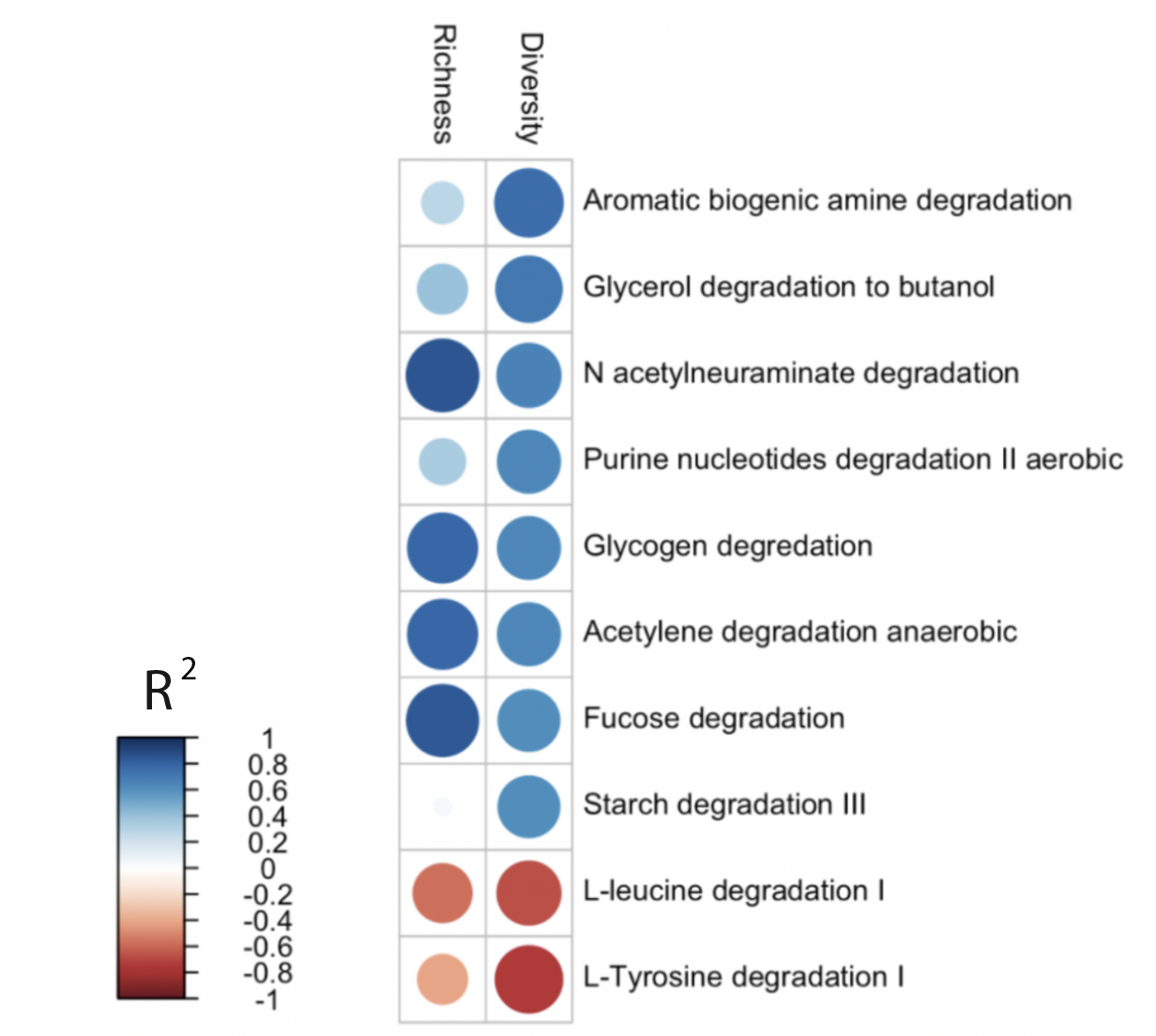

Supplemental Figure 4. Permutational correlations between degradation functional genes identified using PICRUST2 and diversity and richness of land cover (STURLA classes). Strong correlations are set at a threshold of |0.70| with Bonferroni corrections (no significant relationships found).
